# Mapping potential for human encounters with dangerous large mammals across a community-led Wildlife Management Area in southern Tanzania based on overlapping signs of recent activity

**DOI:** 10.64898/2026.10.06.750639

**Authors:** Jude O’Regan, Lucia J. Tarimo, Deogratius R. Kavishe, Fidelma Butler, Aristides Litaka, Manase Elisa, Mohamedi Kamuna, Felister Mombo, Ramiro D. Crego, Gerry F. Killeen

## Abstract

This study examined the spatial distribution of potential human encounters with dangerous large mammals in and adjacent to the Ifakara-Lupiro-Mang’ula Wildlife Management Area (ILUMA WMA) and Nyerere National Park (NNP) in southern Tanzania. Using published survey data collected over one year, this secondary analysis focused on four potentially dangerous large mammals: elephant (*Loxodonta africana*), buffalo (*Syncerus caffer*), hippopotamus (*Hippopotamus amphibius*) and hyena (*Crocuta crocuta*). Potential encounter risk was estimated using overlap indices based on co-occurrence of signs of wildlife and human activities. Potential encounter risk was found to be concentrated around readily identifiable foci within the study area and differed between species and distinguishable human livelihood activities. Potential for encountering the two most apparently important dangerous animals, elephant and buffalo, was closely associated with authorized fishing camps within the ILUMA WMA and the ecotone between substantially disturbed and largely intact natural land cover. The greatest potential for encountering hippopotamus occurred largely around two river beds, while hyena encounter potential was more closely associated with established villages and converted lands close to the western boundary of the ILUMA WMA. These new maps of hazard and risk suggest that effective *human-wildlife conflict* (HWC) mitigation in this WMA is unlikely to arise from any stand-alone intervention, but from targeted actions that reduce spatiotemporal overlap between people and wildlife by keeping them apart and/or protecting them against each other. Overall, this study demonstrates that mapping potential encounter risk based on spatiotemporal overlap of indirect activity indicators like tracks, spoor and signs provides a proactive and statistically powerful way to identify where hazardous encounters are most likely to occur, offering considerable advantages over retrospective assessments of much sparser observations of actual realised conflict events after they have already occurred. This new spatial risk mapping approach provides a framework for shifting management from reactive responses after conflict occurs towards proactive prevention.

## 1. Introduction

The limitations of exclusionary “fortress conservation” approaches (1–3) have contributed to the development of Community Based Natural Resource Management (CBNRM) strategies and tactics across much of Africa (4–6), including Tanzania. The overall principle of CBNRM is to establish institutional frameworks for devolved governance that enable local stakeholder communities to manage land, ecosystems and biodiversity on a sustainable basis, incentivizing them to do so by allowing them to directly benefit from natural resources in terms of livelihoods, services and infrastructure (7–9). By design, CBNRM places humans and wildlife within shared, dynamic, working landscapes where conservation and livelihood activities are reconciled through decentralized management by local stakeholder communities(10–12).

When people and wildlife coexist in CBNRM, HWC become almost inevitable (13–16). Moreover, often locally-managed CBNRM schemes are deliberately established as buffer zones along the fringes of vertically-managed parks and reserves (17,18), with some of them relying on resident custodian communities living within these devolved conservation areas to perform continuous surveillance functions all year-round (19). Nevertheless, HWC is not an inevitable consequence of increased overlap of humans and wildlife and can be managed on an evidence-informed basis (20). Numerous CBNRM initiatives have demonstrated that biodiversity conservation and sustainable human livelihoods can be successfully integrated, enabling long-term coexistence between people and wildlife (18,19,21,22).

Landscape structure plays an important role in determining where risks of HWC interactions occur (23,24), often resulting in quite a focal distribution of risk. For example, edge effects resulting from forest fragmentation have been shown to increase HWC by increasing the frequency of encounters between people and wildlife as the latter are forced to move through landscapes dominated by the people (23). Also, conflict risk is often concentrated around protected areas, permanent water bodies and ecotones between natural land cover and converted landscapes dominated by human settlement, cultivation and livestock rearing (25–28).

However, the evidence base and methodological options for informing effective management of HWC risk remain limited (20,29). While most studies on HWC risk are conducted retrospectively through interviews and routine records, many relevant cases of death, injury and livelihood loss go unreported because of the constrained monitoring capacities, socioeconomic pressures and legal disincentives that often affect disadvantaged rural stakeholder communities in low-income tropical countries (30). This study therefore reports a secondary analysis of published legacy data from an African CBNRM initiative, specifically a Tanzanian Wildlife Management Area (WMA) (17,31,32), to identify spatial foci where the activities of humans overlap with those of several potentially dangerous wild mammals and then examine how landscape characteristics and spatial structure influence the distribution of potential risk.

## 2. Methods

### 2.1 Study area

This study is a secondary analysis of published legacy data from a longitudinal series of rolling cross sectional surveys of the activities of humans, livestock and wild animals across the Ifakara-Lupiro-Mang’ula (ILUMA) WMA in southern Tanzania, the villages immediately to the west of it, and parts of Nyerere National Park (NNP) to the east of it (Fig 1) (33–38).

**Fig 1.**
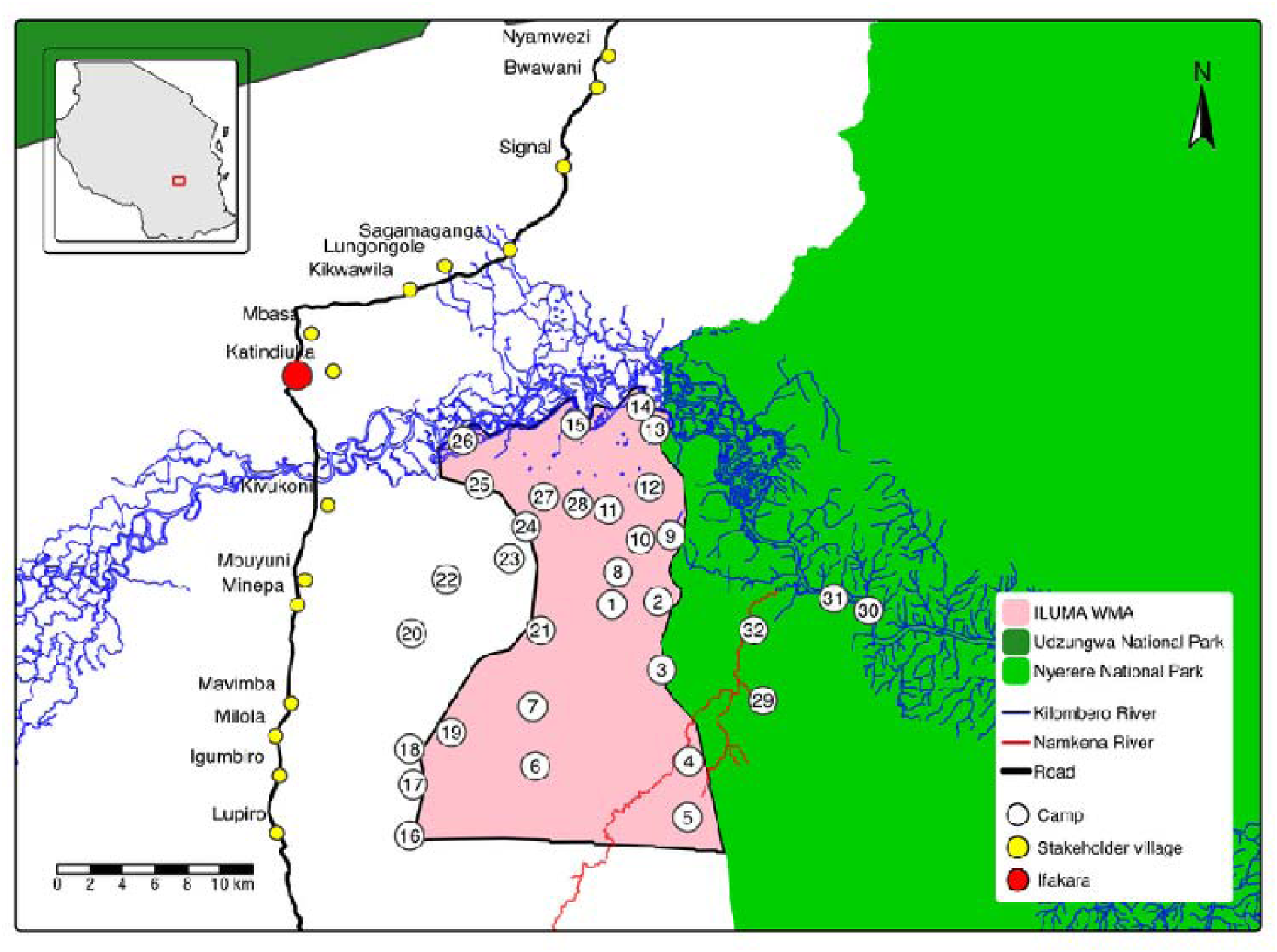
Map of study area showing the location of the 32 survey locations (camps) in and adjacent to ILUMA WMA and inside NNP, within in their local and national conservation context in Tanzania.

The ILUMA WMA (509 km^2^) plays an important role in maintaining connectivity between three major protected areas, namely NNP (formerly part of the Selous Game Reserve), the Kilombero Game Reserve and Udzungwa Mountains National Park (Fig 1). The study area lies on the eastern periphery of the Kilombero Valley, one of the largest freshwater wetlands in East Africa, encompassing a diversity of land cover and usage types, ranging from well-established villages immediately to the west of the WMA, to fully protected land well inside NNP to the east. The natural land cover of the ILUMA WMA is a mosaic of groundwater forests, miombo woodlands, and seasonally flooded grasslands, whereas the surveyed area of NNP immediately to the east of the study area is characterised by drier acacia savanna. The Kilombero river runs along the northern boundary of the WMA, where it supports a number of WMA-authorized and regulated fishing camps established along its banks within the WMA (19,34). The WMA supports a diverse community of wild mammals (37), including large and potentially dangerous herbivores like elephant, buffalo and hippopotamus, all of which have become increasingly common within the WMA as wildlife populations have recovered over recent years. Potentially dangerous large carnivores in the area include lion (*Panthera leo*), leopard (*Panthera pardus*) and spotted hyena (37).

### 2.2 Survey design

The data analysed in this study were collected in 2022 as part of a broader ongoing project investigating malaria vector mosquito ecology. As a result, the original fieldwork focused radial surveys (37) of human, livestock and wild mammal activities around perennial sources of surface water (35,36). Correspondingly, survey locations were purposely selected along rivers, waterholes and stream beds (19,33,34,37). While these selected survey locations were initially distributed across the ILUMA WMA and neighbouring village lands along its western boundary, they were later extended eastwards into NNP to capture less disturbed ecosystems. The survey locations were selected to encompass a wide range of ecosystems, spanning the WMA, legally established villages immediately to the west of it and essentially intact natural ecosystems in NNP to the east.

A total of three survey rounds were conducted across 32 locations, each referred to as a *camp* because the team camped overnight at each of them. Round 1 was conducted between January and February, round 2 between March and May and round 3 between August and November 2022. Rounds 1 and 2 were conducted during the wet season and round 3 took place during the dry season. Survey locations remained at approximately fixed GPS coordinates throughout all rounds, allowing direct comparison of wildlife and human activity across habitat types and disturbance gradients across all three rounds. Each camp was assigned to one of five survey circuits, with each circuit comprising of a set of camps surveyed together. Each of the four circuits within the WMA started and ended at *Msakamba* at the centre of the WMA (Camp 1 in Fig 1), broadly representing the southeast, southwest, northeast and northwest quarters of the ILUMA WMA (37). The fifth circuit, only surveyed in the third round, consisted of 4 camps (camps 29 to 32 in Fig 1) immediately to the east of the WMA, just inside NNP.

At each survey location, radial surveys were conducted by inspecting the periphery of nearby water bodies for various tracks, spoor and other signs of animal and human activity (37). First, the periphery of the water body in closest proximity to where the team camped overnight was surveyed, after which each subsequent water body was circumnavigated and surveyed before proceeding to the next, also surveying along the route between water bodies. These radial surveys were conducted to a maximum distance of 2 km or maximum duration of 4 hours, whichever limit was reached first.

All signs of human, livestock, and wildlife activity encountered along the survey route were recorded. Observations included both direct sightings and indirect evidence such as tracks, dung, feeding signs, and other activity indicators, identified by experienced Village Game Scout (VGS) trackers, using standardised coding keys, guides and data entry sheets as described in detail by Duggan *et al.* (37). For the purposes of this analysis, however, data for only six wildlife species were selected because of their known association with fatal or life-altering outcomes of encounters with humans, including severe impacts upon household livelihoods and resilience: elephant, buffalo, hippopotamus, lion, leopard and spotted hyenas (39–41).

### 2.3 Data analysis and mapping

The published data from rounds 1 to 3 (33,34,37) were selected as these survey rounds collectively span all seasons within a single year, providing a temporally concentrated cross-sectional snapshot of human and wildlife activity detections. These rounds represent data collection prior to substantial development of the WMA, allowing patterns of conflict to be assessed under baseline conditions. The data were subset to focus on indicators of animal and human activity relevant to the objectives of this partial secondary analysis. The dataset was further refined to include only recently made (≤ 24 hours old) tracks and signs, as well as direct observations of humans and animals, to ensure reasonably close temporal overlap between the activities of these selected wildlife species and humans. Livelihood-related activities within and around the WMA include farming, livestock herding, charcoal burning, timber harvesting and, in some cases, illegal wildlife hunting. The extent to which these activities are permitted within each WMA varies according to approved village land-use plans, WMA management plans and locally enacted by-laws, with some livelihood activities being accommodated as part of community-based resource use, while other may be restricted or prohibited. In ILUMA WMA, the land-use plan provides for designated fishing camps where fishing and associated livelihood activities are permitted (34), while charcoal burning, timber harvesting, farming and illegal hunting are not permitted. These activities were therefore classified as representing a particularly vulnerable subset of human activity because, where they occur within conserved wildlife habitat, contrary to the land-use provisions, they may require people to spend extended periods in areas where they may encounter dangerous wildlife, often in poorly constructed shelters (Fig 2). However, even the mere presence of a community member (distinguished from rangers, game scouts and other professionals by the prints left by their footwear) inside the WMA conservation area but outside of the authorized fishing camps (19,34) was assumed to be associated with such unauthorized activities and similar living conditions to those illustrated in Fig 2.

**Fig 2.**
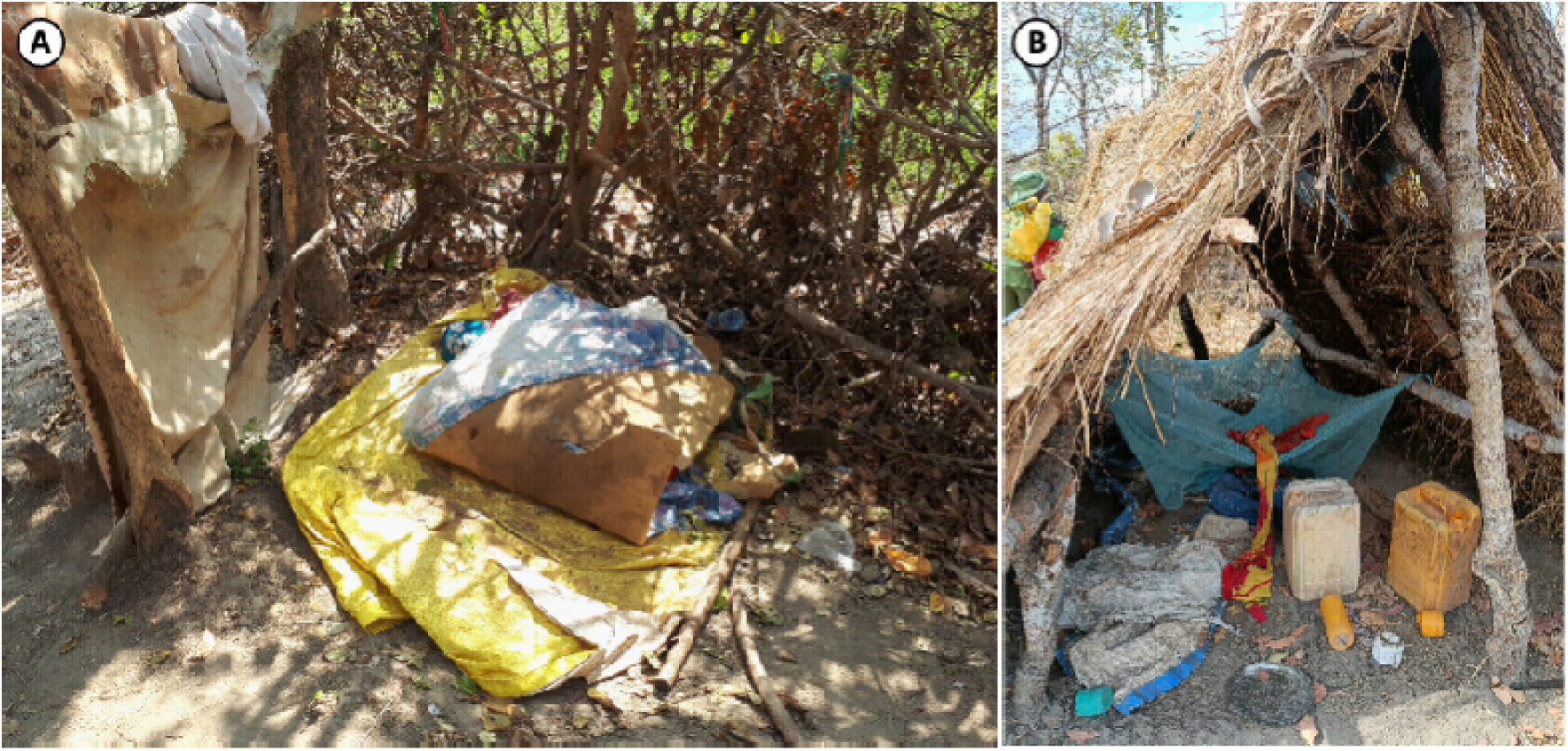
Examples of the living conditions associated with individuals engaged in activities that render them vulnerable to attacks from dangerous wildlife, specifically the temporary camps of an (**A**) unauthorized charcoal burner, and (**B**) an unauthorized rice farmer, both found deep inside the conservation area of the ILUMA WMA (42). While both are clearly vulnerable to attack by predators at night, the latter may also have to defend their nearby crops against large herbivores.

The data were aggregated by survey round to generate total counts of wildlife and human activity detections for each species at each survey location in each round. The summary outcome variables generated reflected the average count per camp in each round of detections of activity by all humans, vulnerable humans, elephant, buffalo, hippopotamus, hyena, leopard and lion. Averages were calculated per survey location (camp) per round because some survey locations were not surveyed in every round. Averages across the three survey rounds eliminates autocorrelation between repeated measures within survey locations.

To estimate potential overlap between humans and wildlife at each survey location, two different human-wildlife activity overlap indices () were calculated for each species. These indices were calculated by multiplying the number of detections of that species (, where *w*= elephant, buffalo, hippopotamus, hyena, leopard or lion) by the number of human activity detections (*N_h_*where *h*= total humans or vulnerable humans):

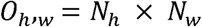

These 12 different indices provide indicators of potential overlap between humans and target species in time and space, where higher values indicate greater potential for dangerous encounters. In the context of the classical risk management literature (43), indices based on overlap with all human activities may be thought of as merely indicators of *hazard*, whereas those based on overlap with humans exhibiting activities that make them obviously far more vulnerable to the animal in question may be considered more proximal indicators of *risk* per se.

Generalised linear models (GLM) were fitted in *R* using the *glmmTMB* (*44*) package to investigate spatial and land-use variables as potential predictors of variation in these human-wildlife overlap indices across the study area. A forward stepwise selection procedure was used to parsimoniously optimize the final fitted models for each dependent variable, with independent variables prioritized for inclusion and retention based on a threshold value of P ≤ 0.05 and at least equivalent AIC estimates to the same model with that variable excluded.

Dependent variables included overlap indices for each of the two indicators of human activity (*N_h_*) and each of the six potentially dangerous wildlife species (*N_w_*). Because the dependent variables were highly skewed and overdispersed, several candidate distributions for these outcomes were tested for optimal goodness of fit, including Poisson and negative binomial distributions, each with and without zero inflation based on the Akaike Information Criterion (AIC) values obtained. Independent variables tested included both a categorical land-use variable and continuous distance variables.

Each radial survey location was classified subjectively into four land-use categories based on the settlement, land use and land cover characteristics described in Kavishe *et al.* (*38*): (1) Established villages to the west of the WMA conservation area, (2) Peripheral settlements inside and to the west of the WMA, (3) Authorized fishing camps established along the banks of the Kilombero river in the north of the WMA (19,34) and largely intact natural areas. For the purposes of mapping and assessing the influence of ecotones, these categories were then grouped into two broader categories: (1) Disturbed areas, comprising outside villages combined with peripheral settlements and (2) Predominantly natural areas, combining fishing camps with intact natural areas.

The geographic distributions of relevant dependent and independent variables were cartographically represented using the *tmap* package of *R*, with the following distance variables calculated using the *sf* (45) package: (1) Distance to either the nearest camp of the other binary land cover category (Disturbed areas versus predominantly natural areas, as per Kavishe *et al.* (38) and explained above) or NNP border (km), whichever was closest, (2) Distance to nearest riverbed (km), and (3) Distance to the nearest western boundary of ILUMA WMA (km).

Distance to either the nearest camp of the other binary land cover category was calculated as the distance to the nearest survey location of the opposite category for each disturbed (villages to the west and peripheral settlements) and natural (fishing camps and natural areas) area. However, with respect to camps 4 and 5 (both peripheral settlements), the boundary of NNP was treated as an equivalent interface between natural landcover and disturbed areas. Following exploratory mapping of elephant and buffalo records, distance to the nearest camp of other category or NNP boundary was included to represent the potential influence of habitat edges and ecotones on wildlife activity and human exposure, as transitional zones have been shown to influence wildlife movement and encounter rates (46,47). In order to facilitate the modelling analysis described below, a modulus transformation was applied to this estimated distance to nearest survey location in the other binary land cover category, as the initial form of this variable contained both negative and positive values, depending on whether that location was respectively on the natural or disturbed side of the ecotone.

Distance to the nearest riverbed, specifically either the Kilombero river or the Namkena river, was included because water availability represents an important ecological driver of wildlife distribution, particularly hippopotamus that need aquatic habitats of sufficient depth (48). Rivers also represent important foci for human activity within the WMA, particularly fishing (19,34) and transport, creating potential overlap between human and wildlife resource use.

Distance to the western boundary of the WMA was included to represent proximity to areas dominated by larger established villages and surrounding land that had largely been converted to farmland, with correspondingly high levels of legal human activity. This independent variable was particularly relevant to hyena, for which exploratory mapping indicated greatest overlap with villages and peripheral settlements in the western portion of the study area.

Two simple transformations of each of these distance variables were also assessed as a means to improve model fit: The untransformed variable, the base 10 logarithm and square root of each distance variable were tested as alternative independent variables in models of each dependent variable, with the form yielding the lowest AIC used for subsequent model selection.

All analyses were conducted using *R*.

## 3. Results

### 3.1 Overall distributions of potentially dangerous animals and their overlaps with people

The total number of detections differed substantially between species, with elephant accounting for almost half of all detections (Fig 3). Buffalo represented the second most frequently recorded species of dangerous large mammal, accounting for over a third of all detections, while hippopotamus and hyena were far less commonly detected. Leopard and lion were only detected 7 and 6 times, respectively, so they were excluded from further analysis.

**Fig 3.**
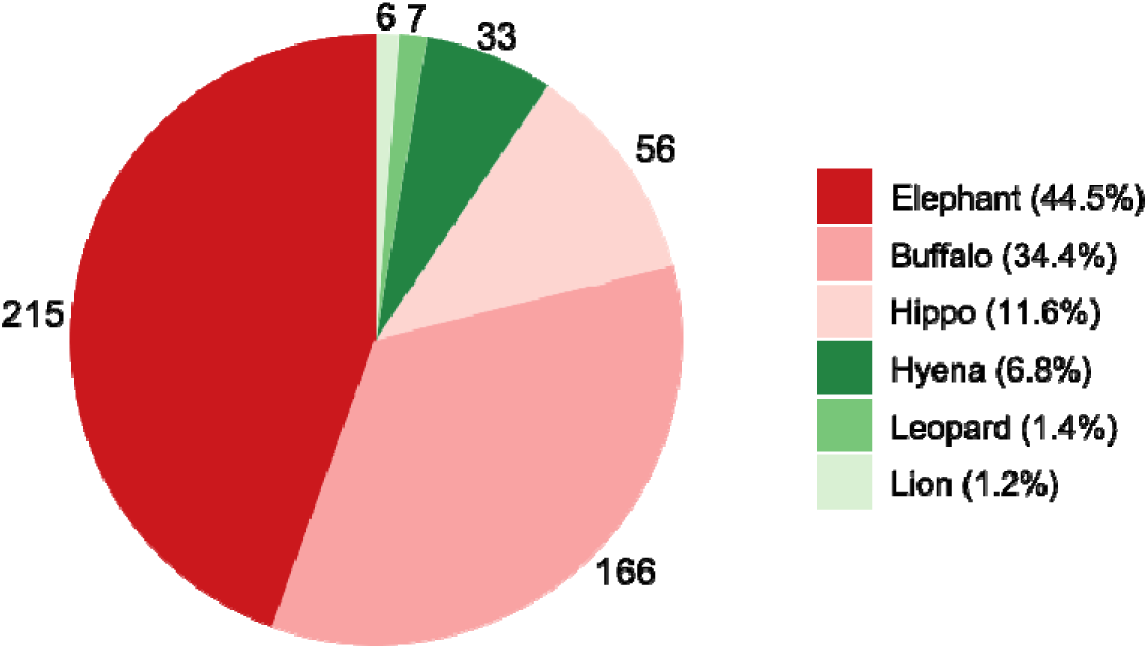
Total detections of the six species of potentially dangerous large mammals across the study area, summed across all three survey rounds.

Overall, the activity distributions of elephant, buffalo, hippopotamus and hyena were highly focal (Fig 4, panels C to F) and, with some important exceptions, largely occurred in different geographic areas to those of humans (Fig 4, panels A and B). Highest human activity was concentrated in the west and southwest of the WMA (Fig 4, panel A), in and around the peripheral settlements and villages, whereas elephant, buffalo and hippopotamus activity was concentrated mainly in the east and north, within the natural areas along the river and floodplain margin (Fig 4, panels C, D and E). Hyena activity broke from this pattern, spreading further west into areas closer to the peripheral settlements and villages. Notably, most of the human activity recorded inside the WMA boundary was classified as vulnerable human activity (Fig 4, panel B), indicating that the bulk of human presence within the reserve itself consisted of unauthorized or high-risk activities rather than general human activity.

**Fig 4.**
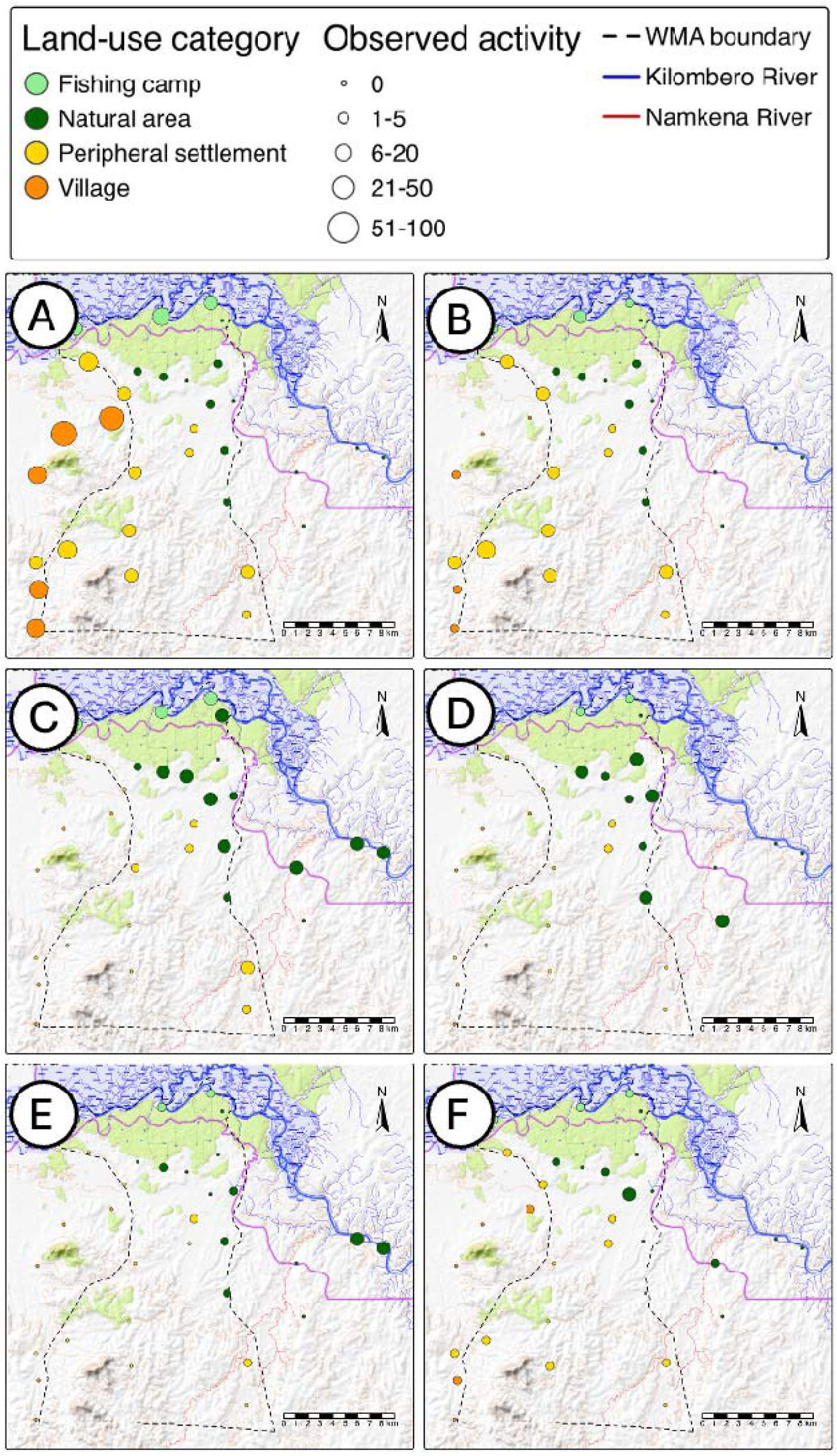
Spatial distribution of mean activity for (**A**) all humans, (**B**) vulnerable humans, (**C**) elephant, (**D**) buffalo, (**E**) hippopotamus and (**F**) hyena across the study area.

Correspondingly, the frequency distributions of activity detection overlaps between potentially dangerous large mammals and all humans (Fig 5A), or vulnerable humans specifically (Fig 5B), respectively indicates that elephant probably represent the greatest overall hazard and risk. In terms of contributions to hazard and risk, elephant are followed by buffalo, hippopotamus and hyena, while leopard and lion exhibited almost no overlap with people. For all four of the other large, potentially dangerous animals, the distributions of overlap indices were highly overdispersed, indicating highly focal concentration of hazard and risk in a small minority of locations: most overlaps occurring at relatively low values for the relevant index, while a remarkably small fraction of times and places exhibited unusually high activity levels for both humans and one or more dangerous wild animals (Fig 5). While over three quarters of all observations recorded no overlap of any humans with any of these wild animals, three quarters of the total estimated overlap across species was accounted for by 16% or less of all observations (Fig 5).

**Fig 5.**
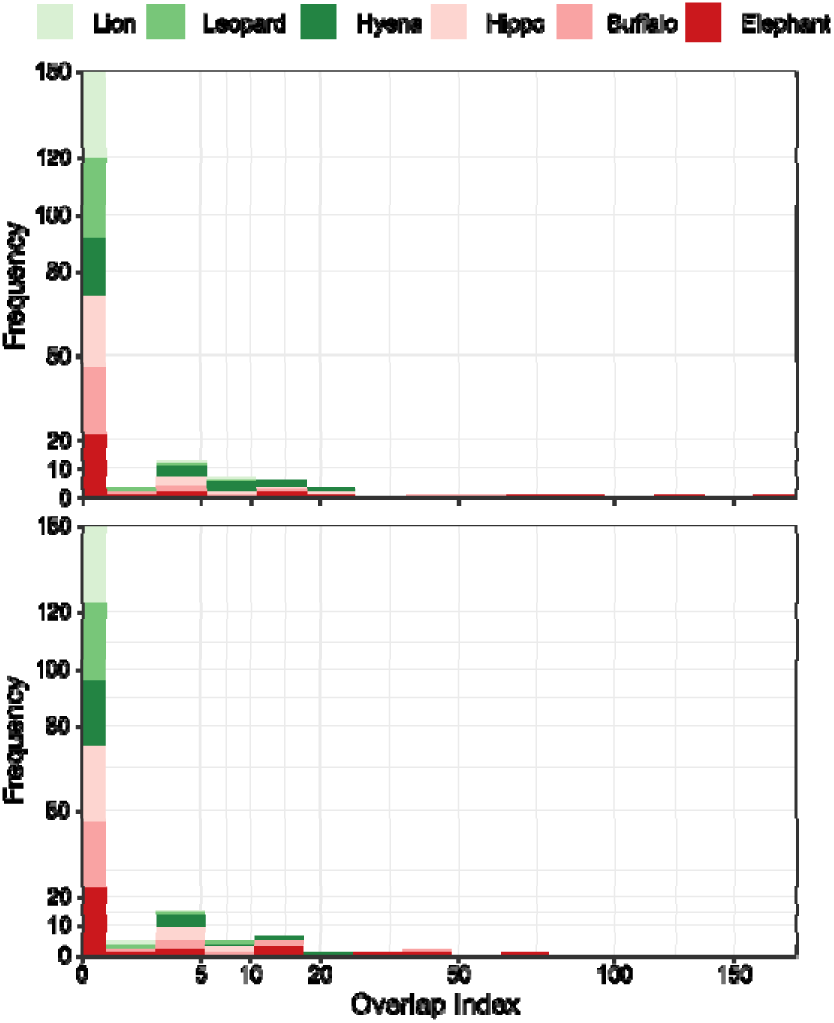
Frequency distributions of the overlap indices for 6 potentially dangerous wild mammals across the area with (**A**) all humans and (**B**) vulnerable humans only.

### 3.2 Activity overlaps between humans and elephant

Elephant overlap with humans in general is unevenly distributed across the study area, with the authorised fishing camps established amidst the groundwater forest along the south bank of the Kilombero river exhibiting the greatest apparent potential for hazardous encounters (Fig 6A). Most other overlaps with all humans occur near the boundary between natural areas and peripheral settlements, indicating that hazard is particularly associated with the land cover ecotone, where human-occupied lands that are at least partially converted meets relatively intact natural elephant habitat. While villages outside of the conservation area exhibited absolutely no signs of overlap between elephant and any humans, there was also little evidence for such hazards anywhere inside NNP, or outside of fishing camps in the best conserved areas in the northeast of ILUMA.

**Fig 6.**
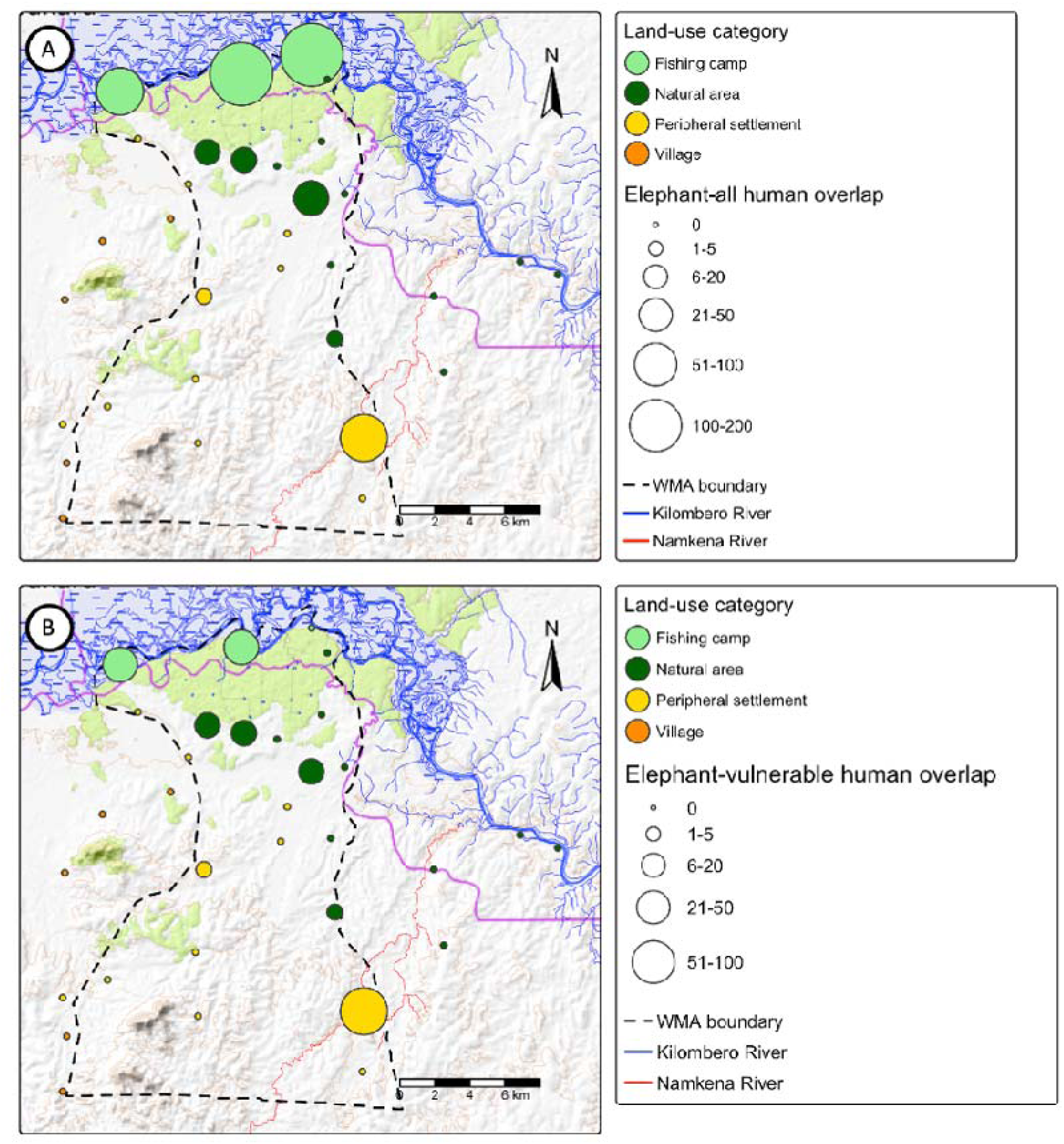
Spatial distribution of mean indices for elephant activity overlap across the study area with either (**A**) all humans or (**B**) vulnerable humans specifically.

However, focusing specifically upon specific detected activities of humans likely to be especially vulnerable to hazardous encounters with these large mammals (Fig 6B) reveals a generally similar spatial pattern but with a much lower fraction of overall activity overlap attributed to the fishing camps, especially the easternmost of the three along the NNP boundary. Indeed, for those people who stay inside the WMA under the most vulnerable of circumstances because they do so without permission (Fig 2), such risks appear particularly closely associated with proximity to that ecotone.

GLM analysis confirms that, relative to fishing camps, activity overlap between elephant and all humans was dramatically lower in all other land use categories (Table 1): overlap hazard was estimated to be 97% lower in predominantly natural areas, 98% lower in peripheral settlements, and so low in villages outside the WMA that it was not estimable by GLM and therefore essentially negligible. Also, modulus of distance to the nearest camp of the opposite disturbed versus natural stratum or the NNP boundary, representing the ecotone between intact natural land cover and substantially human-modified landscapes, exhibited a negative trend with increasing distance that approached significance (Table 1 and Fig 7).

**Fig 7.**
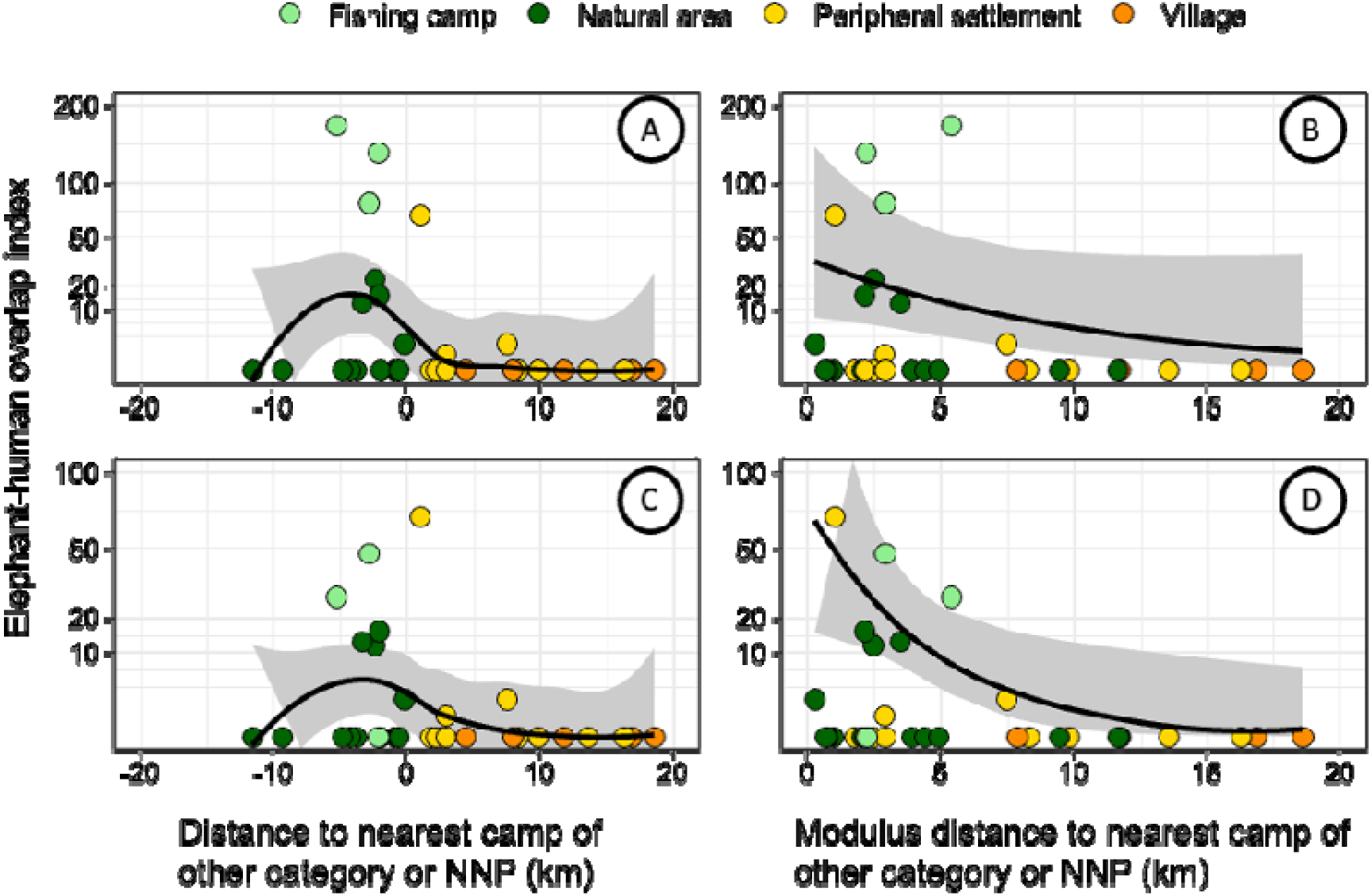
Relationship between distance and modulus of distance to the nearest camp of the other category (disturbed areas versus essentially intact natural areas) or the Nyerere National Park (NNP) boundary and the elephant–human overlap index. Panels A and C show the relationships using the raw distance predictor (km) for all humans and vulnerable humans, respectively. Panels B and D show the fitted relationships using the modulus-transformed distance predictor for all humans and vulnerable humans, respectively. In panels A and C, the solid line represents a fitted LOESS smooth and the shaded ribbon represents the derived 95% confidence interval. In panels B and D, the solid line represents the prediction of the fitted model in table 1 and the shaded ribbon represents the estimated 95% confidence interval for that model fit.

**Table 1.** Results of negative binomial general linear models examining the effect of land use category and modulus of the distance to nearest camp of the other category or NNP (representing distance to the ecotone of natural land cover and converted land cover) on the human-wildlife overlap indices for elephant across the study area.

| Dependent and independent variables | $\beta \pm \text{SEM}$ | RR [95% CI] | P |
| --- | --- | --- | --- |
| <u>Elephant activity overlap with all humans (Negative binomial type 1)</u> |  |  |  |
| *Land use category* , AIC = 117.7 |  |  |  |
| Fishing camp | NA | NA | NA |
| **Natural area** | **-3.49 ± 0.66** | **0.03 [0.01, 0.11]** | **<0.001** |
| **Peripheral settlement** | **-4.11 ± 0.84** | **0.02 [0.00, 0.08]** | **<0.001** |
| Village | -23.12 ± NE | NE | NE |
| *Modulus distance to nearest camp of other category or NNP border (per km)*, AIC =130.3 | -0.19 ± 0.11 | 0.82 [0.66, 1.02] | 0.083 |
| <u>Elephant with vulnerable humans (zero-inflated negative binomial type 2)</u> |  |  |  |
| *Land use category*, AIC = 108.0 |  |  |  |
| Fishing camp | NA | NA | NA |
| Natural area | -1.31 ± 0.78 | 0.27 [0.06, 1.23] | 0.090 |
| Peripheral settlement | -0.09 ± 0.88 | 0.91 [0.16, 5.14] | 0.919 |
| Village | -23.15 ± NE | NE | NE |
| *Modulus distance to nearest camp of other category or NNP border (per km)*, AIC = 106.1 | -0.43 ± 0.19 | 0.65 [0.45, 0.94] | 0.022 |

CI: Confidence interval RR: Relative rate NA: Not applicable because reference group NE: Parameter not estimable because no overlapping detections recorded resulting in unstable model estimates so excluded from final model SEM: Standard error of the mean

For vulnerable people specifically, GLM analysis indicated that encounter risk in natural areas appeared to be only one quarter of that in fishing camps, although that contrast only approached significance (Table 1). However, peripheral settlements appeared to be just as hazardous as fishing camps (Table 1). Established villages to the west of the WMA exhibited essentially no overlap between elephant and vulnerable humans, so encounter risk appeared to be essentially negligible. In contrast, risk of overlap was closely associated with proximity to the ecotone between essentially intact natural land cover and substantially disturbed lands, declining by approximately 35% per kilometre further away from that interface (Table 1 and Fig 7).

Regarding possible encounters between elephant and all humans, the greatest hazards occur closest to the ecotone but is nevertheless highest among the 3 fishing camps, even the one furthest from the ecotone (Figs 6 and 7). Although only approaching significance in Table 1, elephant encounter hazard for all humans appears to decline sharply with distance from the ecotone and reach essentially zero at approximately 7 km on either side of the ecotone (Fig 7). Regarding activity overlap between vulnerable humans and elephant, a similar but significant trend is observed (Table 1), with all encounter risk occurring within approximately 7 km of the ecotone (Fig 7B).

### 3.3 Activity overlaps between humans and buffalo

The activities of buffalo overlapped with those of humans in a highly focal manner (Fig 8). Regarding buffalo activity overlap with that of humans in general (Fig 8A), the fishing camp *Mdalangwila* (Camp 15 in Fig 1), exhibited the highest hazard, presumably because several pathways through the groundwater forest there are equally popular with both species. All other overlap between buffalo and humans in general is spatially concentrated in natural areas and in two peripheral settlements along the ecotone between disturbed and natural strata (Fig 8A). Similar to total human activity overlap with elephant (Fig 7A), hazard is particularly associated with this interface between human-settled areas and relatively intact natural buffalo habitat occurs. For vulnerable humans specifically (Fig 8B), the spatial distribution of overlap with buffalo was very similar to that for all humans, although encounter risk at *Mdalangwila* was notably lower. Most remaining overlap with vulnerable humans was similarly concentrated in natural areas and peripheral settlements close to the disturbed-natural ecotone.

**Fig 8.**
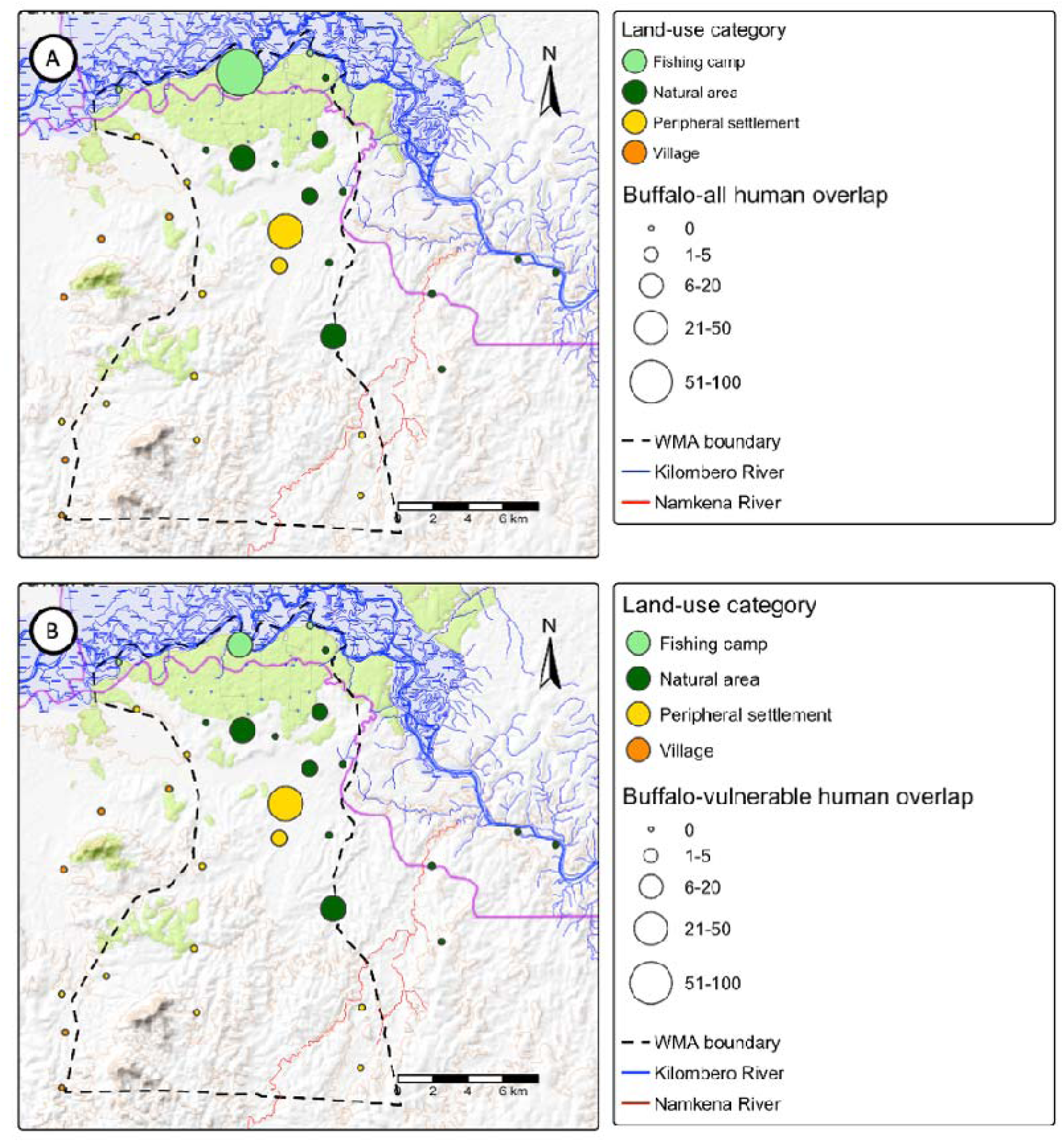
Spatial distribution of mean indices for buffalo activity overlap across the study area with either (**A**) all humans or (**B**) vulnerable humans specifically.

GLM analysis confirms that fishing camps clearly exhibited the greatest levels of overlap between buffalo and all humans, with other natural areas and peripheral settlements respectively estimated to have 86% and 73% lower overlap probability, while encounter hazard in established villages outside the western boundary of the WMA was negligible due to a complete lack of buffalo detections (Table 2). Indeed, besides those in close proximity to the ecotone between disturbed and natural areas, all other camps exhibited no activity overlap between buffalo and humans in general. However, although modulus of distance to the nearest camp of the opposite land cover stratum (disturbed versus predominantly natural) or the NNP boundary appeared to exhibit a negative trend with increasing distance, this relationship was not statistically significant (Table 2).

**Table 2.**
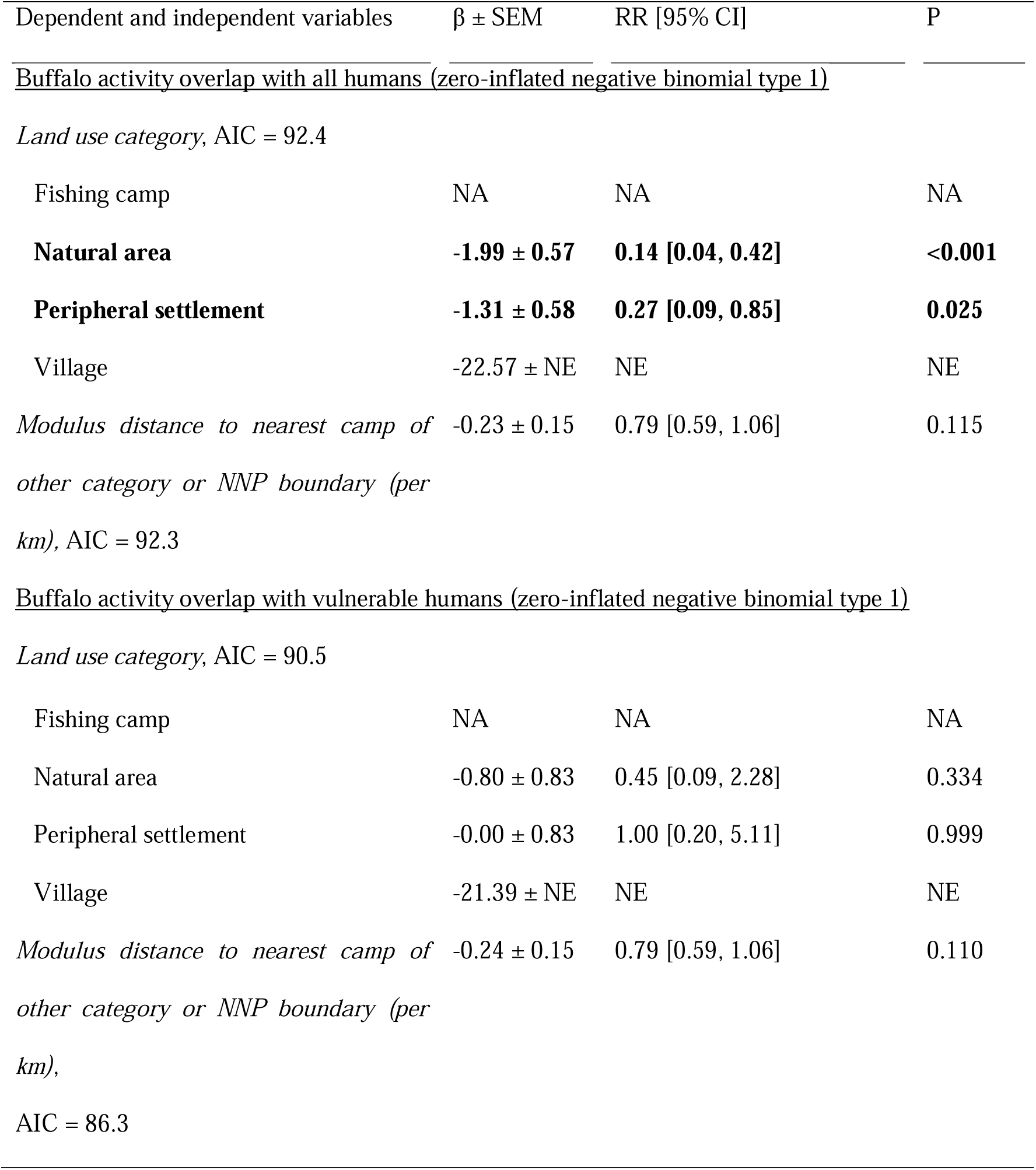
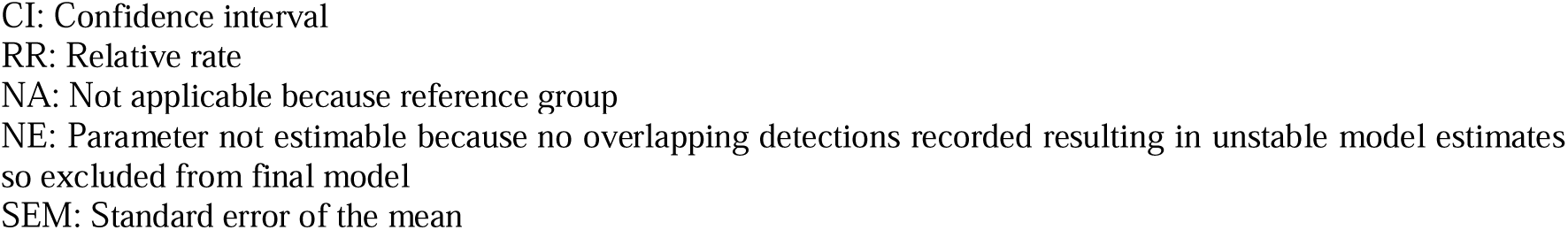
Results of negative binomial generalised linear models examining the effect of land use category and modulus of the distance to nearest camp of the other category (representing distance to the ecotone of natural land cover and converted land cover) on the risk of potentially dangerous encounters between humans and buffalo across the study area.

| Dependent and independent variables | $\beta \pm \text{SEM}$ | RR [95% CI] | P |
| --- | --- | --- | --- |
| <u>Buffalo activity overlap with all humans (zero-inflated negative binomial type 1)</u> |  |  |  |
| *Land use category, AIC = 92.4* |  |  |  |
| Fishing camp | NA | NA | NA |
| **Natural area** | **-1.99 ± 0.57** | **0.14 [0.04, 0.42]** | **<0.001** |
| **Peripheral settlement** | **-1.31 ± 0.58** | **0.27 [0.09, 0.85]** | **0.025** |
| Village | -22.57 ± NE | NE | NE |
| *Modulus distance to nearest camp of other category or NNP boundary (per km), AIC = 92.3* | -0.23 ± 0.15 | 0.79 [0.59, 1.06] | 0.115 |
| <u>Buffalo activity overlap with vulnerable humans (zero-inflated negative binomial type 1)</u> |  |  |  |
| *Land use category, AIC = 90.5* |  |  |  |
| Fishing camp | NA | NA | NA |
| Natural area | -0.80 ± 0.83 | 0.45 [0.09, 2.28] | 0.334 |
| Peripheral settlement | -0.00 ± 0.83 | 1.00 [0.20, 5.11] | 0.999 |
| Village | -21.39 ± NE | NE | NE |
| *Modulus distance to nearest camp of other category or NNP boundary (per km), AIC = 86.3* | -0.24 ± 0.15 | 0.79 [0.59, 1.06] | 0.110 |

CI: Confidence interval RR: Relative rate NA: Not applicable because reference group NE: Parameter not estimable because no overlapping detections recorded resulting in unstable model estimates so excluded from final model SEM: Standard error of the mean

In contrast, overlap between buffalo and vulnerable humans did not differ significantly among fishing camps, natural areas and peripheral settlements, although this may be attributed to highly uncertain model parameter estimates and limited statistical power (Table 2). Similarly to all humans, the buffalo activity overlap with vulnerable humans specifically appeared to decline with the modulus of distance to the nearest camp of the other category (disturbed versus predominantly natural) or the NNP boundary, although this relationship was also not statistically significant (Table 2). Nevertheless, inspection of Fig 9 indicates that the hazards (Panels A and B) and risks (Panels C and D) of encountering buffalo appear concentrated within approximately 6 km on either side of the disturbed versus natural ecotone, where people are most likely to venturing into intact buffalo habitat and buffalo are most likely to venture out of it.

**Fig 9.**
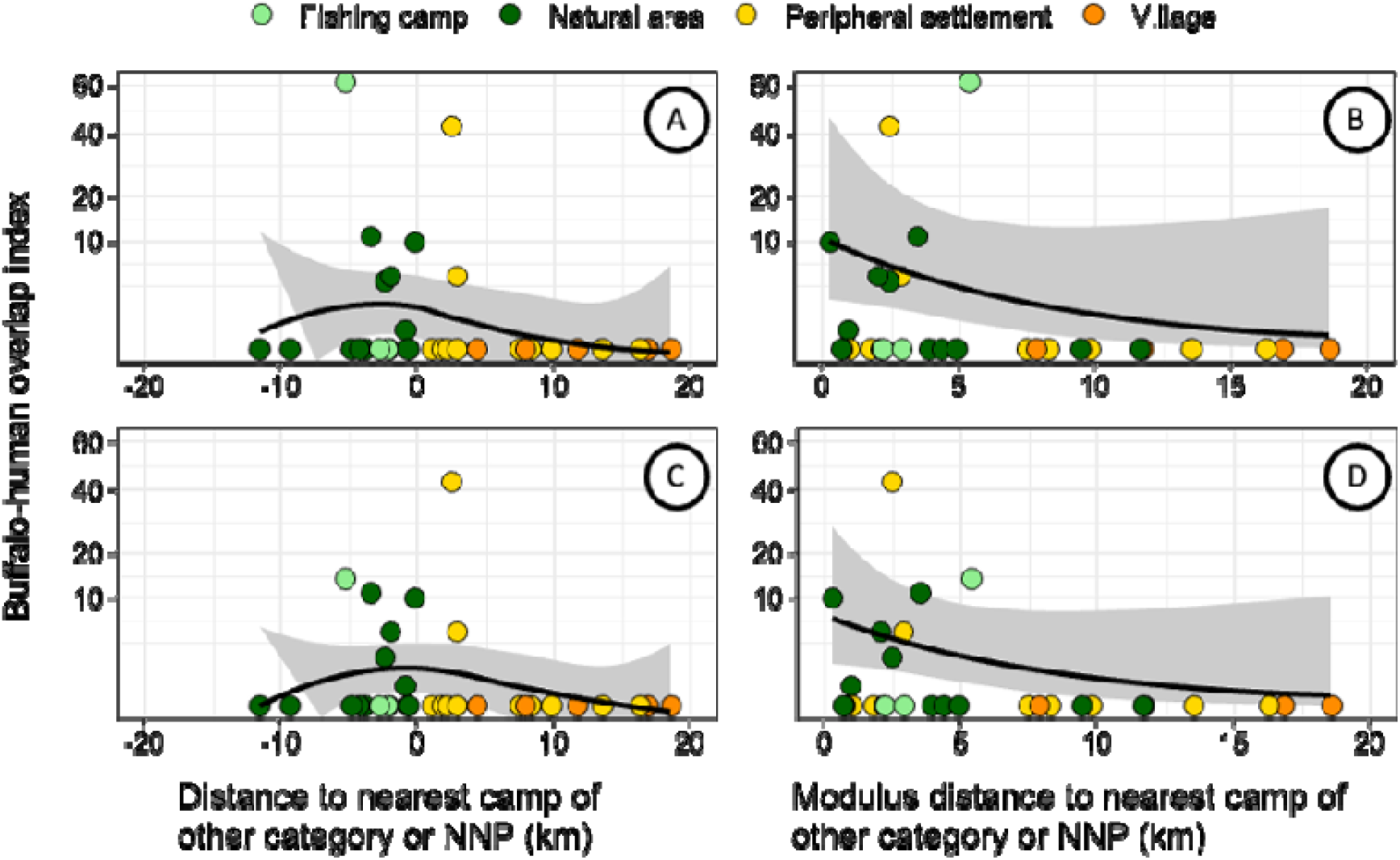
Relationship between distance and modulus of distance to the nearest camp of the other category (disturbed areas versus essentially intact natural areas) or the Nyerere National Park (NNP) boundary and the buffalo–human overlap index. Panels A and C show the relationships using the raw distance predictor (km) for all humans and vulnerable humans, respectively. Panels B and D show the fitted relationships using the modulus-transformed distance predictor for all humans and vulnerable humans, respectively. In panels A and C, the solid line represents a fitted LOESS smooth and the shaded ribbon represents the derived 95% confidence interval. In panels B and D, the solid line represents the prediction of the fitted model in table 2 and the shaded ribbon represents the estimated 95% confidence interval for that model fit.

### 3.4 Activity overlaps between humans and hippopotamus

In contrast, hippopotamus activity overlap with humans is much more highly concentrated in a small number of survey locations within the WMA (Fig 10). For hippopotamus overlap with humans in general (Fig 10A), the fishing camps along the Kilombero river exhibit by far the highest encounter hazard, reflecting the close association between hippopotamus, people and large perennial water sources. Outside these riverside fishing camps, most other surveyed locations exhibited no detectable encounter hazard, the exceptions being two natural areas and one peripheral settlement close to the Namkena river in the southeast, all within the WMA. GLM analysis estimated that natural areas and peripheral settlements have 98% and 99% lower encounter risk than fishing camps, respectively, whereas no hippopotamus activity was detected in the villages outside the WMA (Table 3). Natural areas along the Kilombero River within NNP exhibited no overlap despite frequent hippopotamus detections, reflecting the absence of human activity within these protected areas rather than a lack of hippopotamus. Notably, overlap risk declined significantly with increasing distance from the nearest substantial riverbed, with each unit increase in the square root of distance associated with a 91% reduction in encounter risk (Table 3).

**Fig 10.**
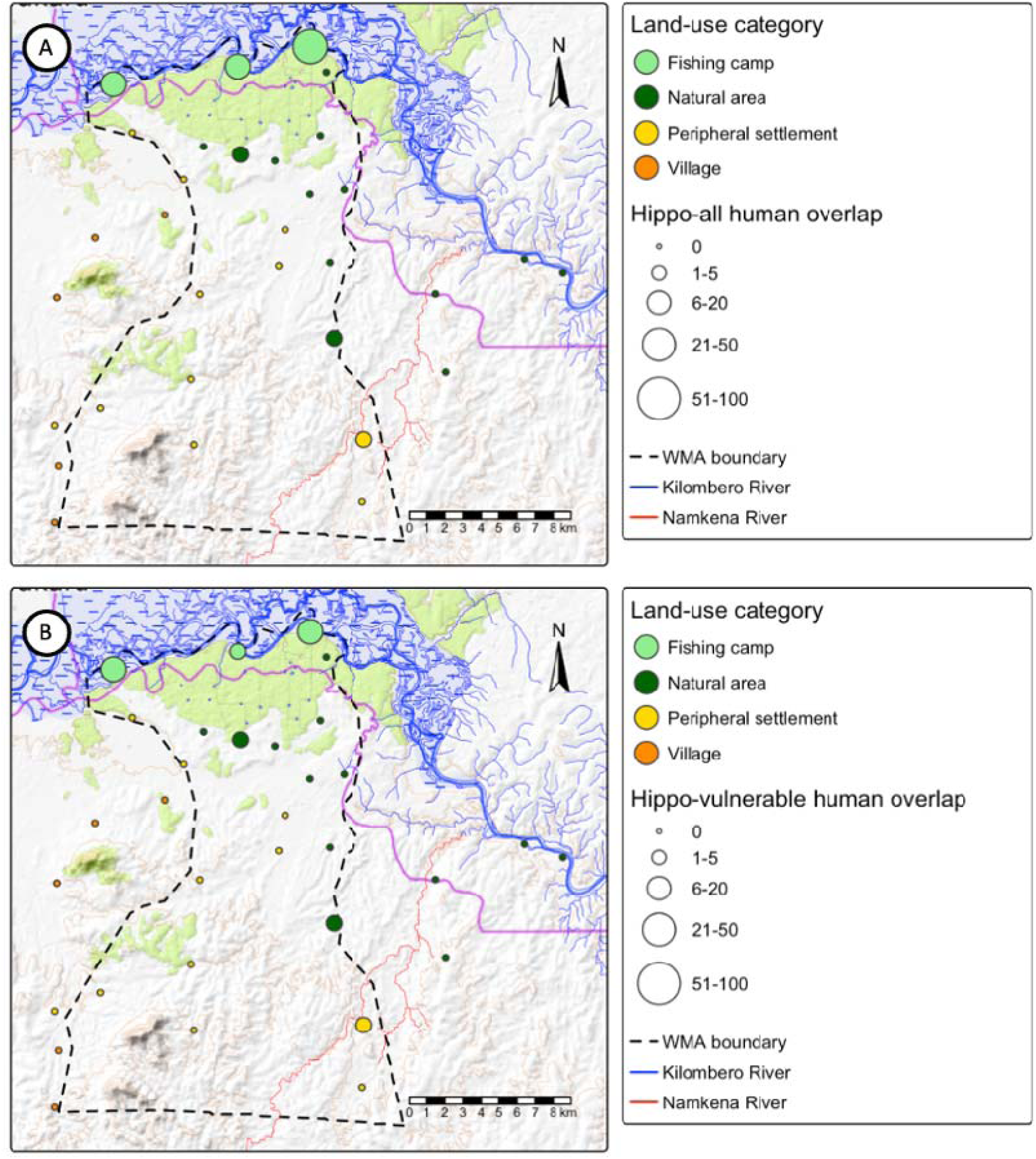
Spatial distribution of mean indices for hippopotamus activity overlap across the study area with either (**A**) all humans or (**B**) vulnerable humans specifically.

**Table 3.**
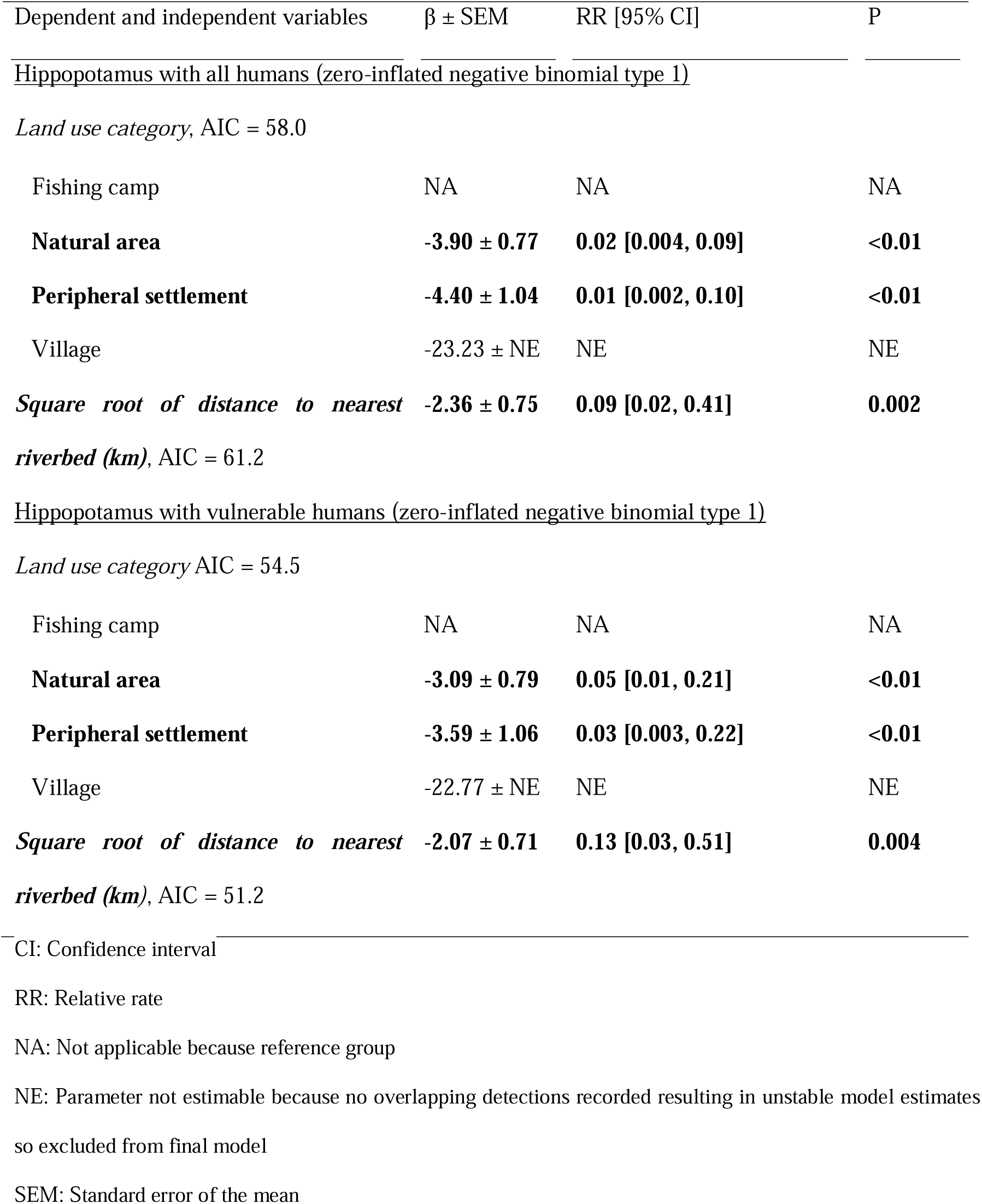
Results of negative binomial general linear models examining the effect of land use category and distance to nearest substantial riverbed on the human-wildlife overlap indices for hippopotamus across the study area.

Regarding risk of hippopotamus encounter for vulnerable humans specifically (Fig 10B), an almost identical spatial pattern was observed, with most overlap occurring at fishing camps. Natural areas and peripheral settlements were estimated to have 95% and 97% lower risk respectively, while villages outside the WMA again exhibited no detections of hippopotamus and therefore negligible risk (Table 3). Similarly, encounter risk decreased significantly with increasing distance from the nearest substantial riverbed, with each unit increase in the square root of distance associated with an 87% reduction in overlap risk (Table 3). Indeed, almost all such interactions with humans in general or with vulnerable humans specifically occurs within 1km of the nearest riverbed and none occurs more than 5 km away from one (Fig 11), reflecting the strong dependence of hippopotamus on substantial water bodies.

**Fig 11.**
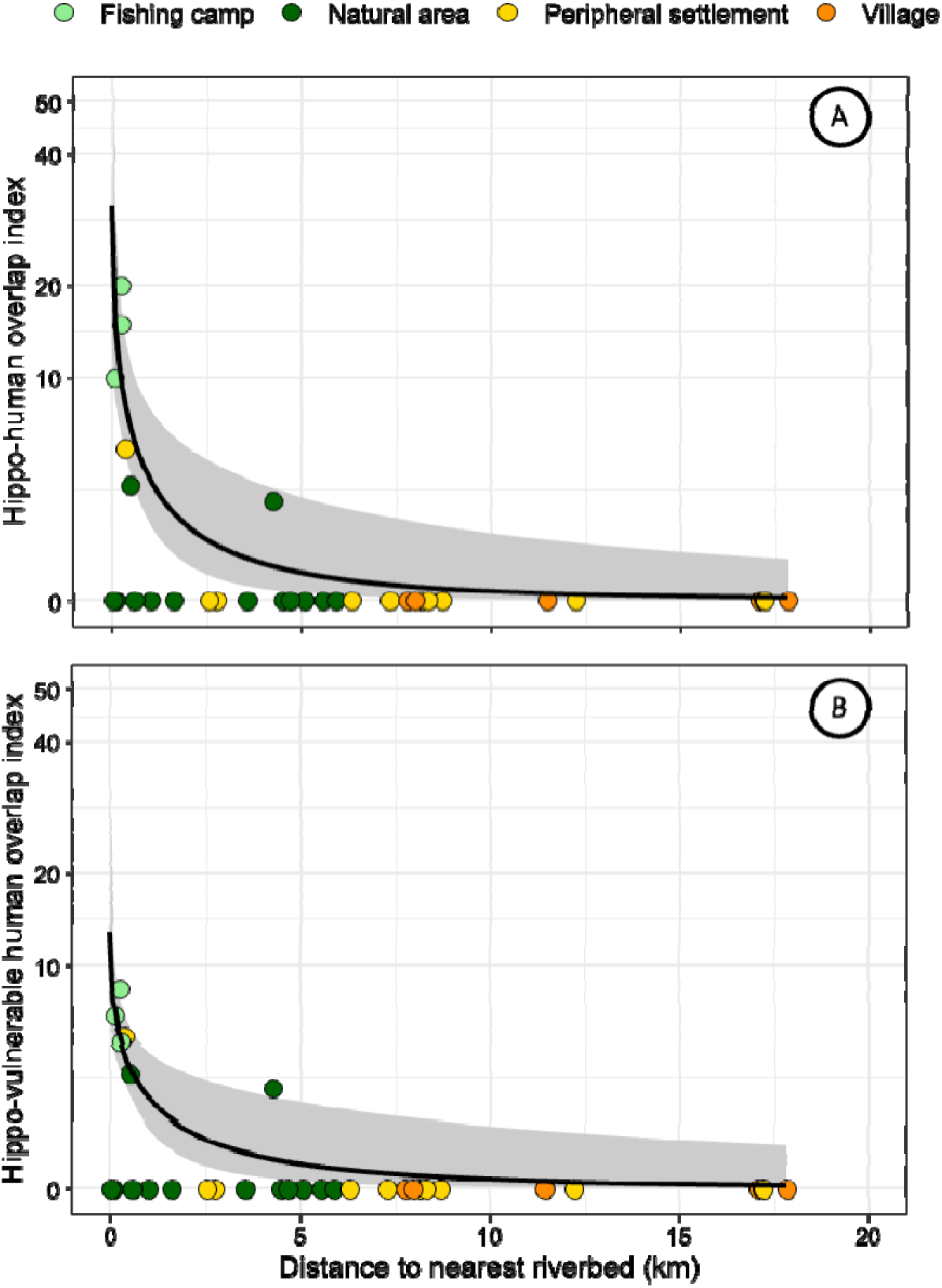
Relationship between distance to the nearest riverbed (km) and the hippopotamus– human overlap index. Panel A shows the relationship for all humans and Panel B shows the relationship for vulnerable humans. In both panels, the solid line represents the prediction of the fitted model in table 3 and the shaded ribbon represents the estimated 95% confidence interval for that model fit.

### 3.5 Activity overlaps between humans and hyena

Surprisingly, risk of human encounters with hyena was concentrated along the western border of the ILUMA WMA, where the landscape becomes dominated by large established villages and peripheral settlements (Fig 12). Hyena activity overlap with humans in general is mainly concentrated around villages and peripheral settlements close to either side of the western and southern borders of the WMA, with all other activity overlap risk being in the two westernmost fishing camps and a small amount in only two other natural areas within the WMA (Fig 12A). Fishing camps again exhibited the highest estimated activity overlap between hyena and humans in general, with natural areas estimated to have almost 90% lower encounter hazard. Although peripheral settlements and villages showed no significant difference in encounter risk relative to fishing camps (Table 4), this arises from considerable variability between such locations and corresponding uncertainty around these estimates that make consistent geographic patterns across those strata difficult to identify. Encounter hazard declined markedly with increasing distance from the western boundary of the ILUMA WMA, more than halving over each unit increase in the square root of distance (Table 4 and Fig 13A). Indeed, almost all hyena encounter hazard occurred within 7 km of the border (Fig 13A).

**Fig 12.**
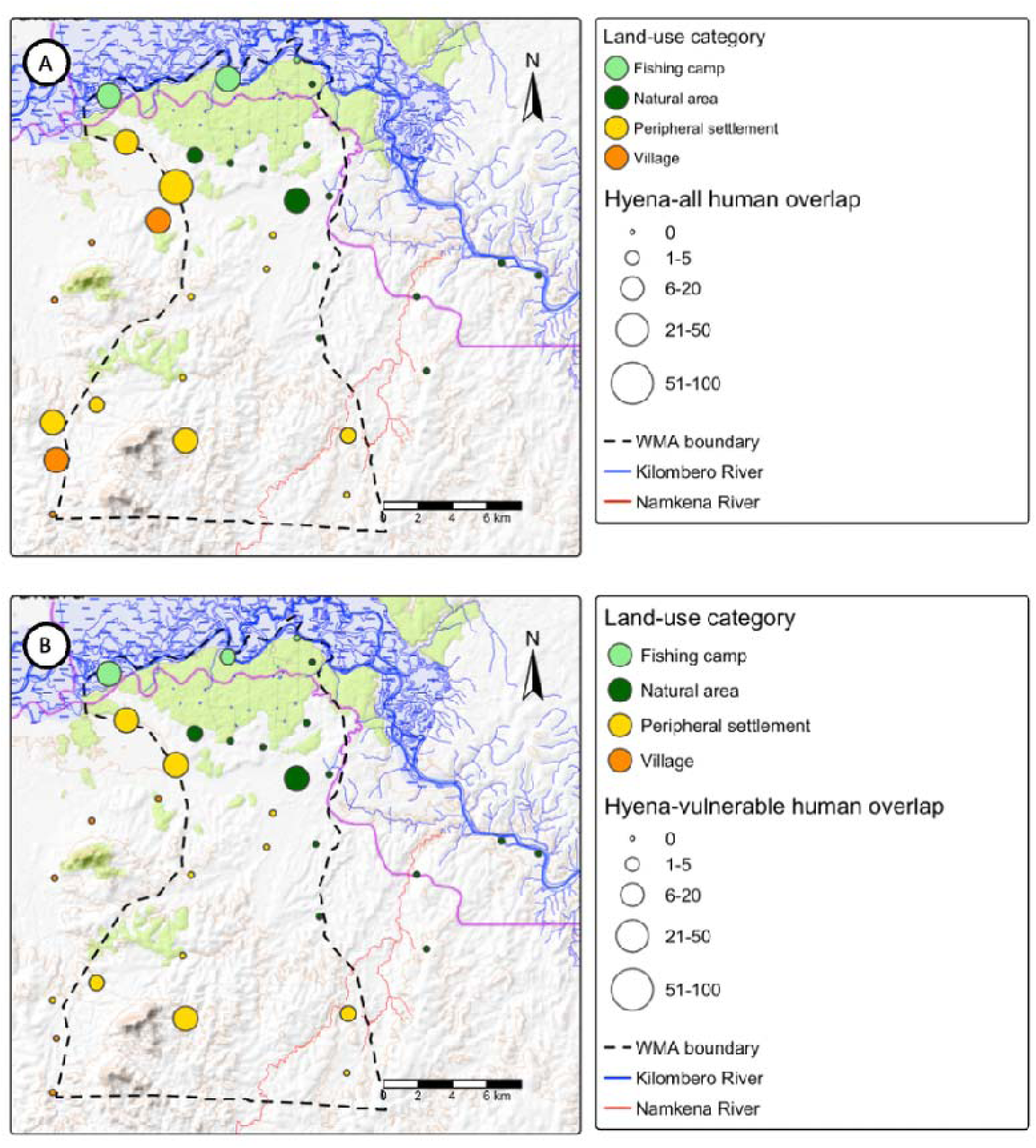
Spatial distribution of mean indices for hyena activity overlap across the study area with either (**A**) all humans or (**B**) vulnerable humans specifically.

**Fig 13.**
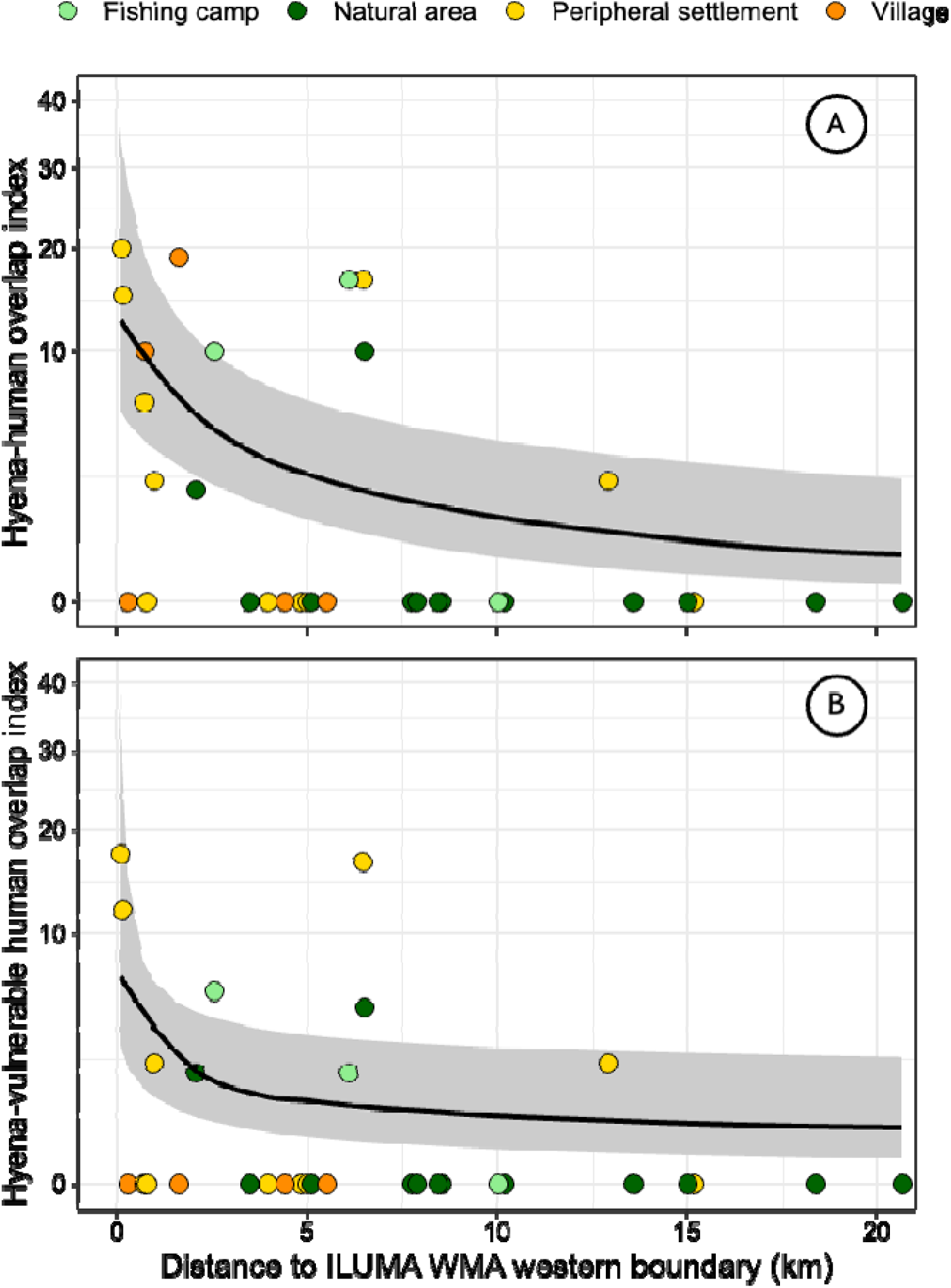
Relationship between distance to the ILUMA WMA western boundary (km) and the hyena–human overlap index. Panel A shows the relationship for all humans and Panel B shows the relationship for vulnerable humans. In both panels, the solid line represents the prediction of the fitted model in table 4 and the shaded ribbon represents the estimated 95% confidence interval for that model fit.

**Table 4.** Results of negative binomial general linear models examining the effect of land use category and distance to the western boundary of the ILUMA WMA on the human-wildlife overlap indices for hyena across the study area.

| Dependent and independent variables | $\beta \pm \text{SEM}$ | RR [95% CI] | P |
| --- | --- | --- | --- |
| <u>Hyena with all humans (Negative binomial type 1)</u> |  |  |  |
| *Land use category, AIC = 131.0* |  |  |  |
| Fishing camp | NA | NA | NA |
| **Natural area** | **-2.17 ± 0.96** | **0.11 [0.02, 0.76]** | **0.025** |
| Peripheral settlement | -0.53 ± 0.76 | 0.59 [0.13, 2.59] | 0.482 |
| Village | -0.83 ± 0.95 | 0.43 [0.07, 2.77] | 0.377 |
| ***Square root of distance to western boundary of ILUMA WMA (km),*** | **-0.88 ± 0.28** | **0.42 [0.24, 0.72]** | **0.002** |
| AIC = 123.2 |  |  |  |
| <u>Hyena with vulnerable humans (Zero-inflated negative binomial type 1)</u> |  |  |  |
| *Land use category, AIC = 93.6* |  |  |  |
| Fishing camp | NA | NA | NA |
| Natural area | -1.77 ± 0.98 | 0.17 [0.03, 1.16] | 0.070 |
| Peripheral settlement | -0.34 ± 0.80 | 0.71 [0.15, 3.42] | 0.668 |
| Village | -20.86 ± NE | NE | NE |
| ***Logarithm of distance to western boundary of ILUMA WMA (km),*** | **-1.27 ± 0.49** | **0.28 [0.11, 0.73]** | **0.009** |
| AIC = 92.7 |  |  |  |

CI Confidence interval RR: Relative rate NA: Not applicable because reference group NE: Parameter not estimable because no overlapping detections recorded resulting in unstable model estimates so excluded from final model SEM: Standard error of the mean

Given the predatory and highly nocturnal nature of hyena, human vulnerability to attack at night (Fig 2) is particularly important, so the distribution of overlap with vulnerable humans specifically is particularly important in this case. Fig 12B reveals that hyena activity overlap with vulnerable humans in particular appears notably absent from villages outside the WMA but occurs most frequently in peripheral settlements just inside western boundary of the WMA, although some overlap also occurs in the same two westernmost fishing villages and two natural areas as for humans in general (Fig 12A). Fishing camps again exhibited the highest estimated mean risk of encounters with hyenas for vulnerable humans but risk was again highly variable within the peripheral settlements. Correspondingly, while GLM analysis indicated that natural areas exhibited >80% less encounter risk than fishing camps, this difference only approached significance because of the associated uncertainty around these estimates. Hyena activity overlaps with vulnerable human activities were not recorded in villages outside the WMA, resulting in encounter risk that could not be estimated for villages by GLM (Table 4). Note, however, that this impression may be somewhat biased by the fact that many such observations may have been misclassified because the detected activities were not classified as being associated with vulnerability, simply because they are illegal within the WMA but often legal outside it. Hyena encounter risk for vulnerable humans noticeably declined with increasing distance from the western boundary of the ILUMA WMA, decreasing by almost three-quarters per unit increase in the logarithm-transformed distance (Table 4 and Fig 13B).

## 4. Discussion

### 4.1 Drivers of risk

Many studies have assessed drivers of HWC, almost always retrospectively, based on interviews, activity-based indicators of pre-conflict behaviours like daytime use of densely vegetated habitat that precede nocturnal incursions into croplands and settlements, or routine records of relevant events, such as human casualties, depredation of livestock or crop damage (49–52). However, this study took the alternative approach of mapping potential for human encounters with dangerous large mammals before they occur and assessing how survey location and landscape characteristics influence those hazards and risks, based on spatial and temporal overlap of largely indirect signs of activity, such as tracks and spoor. Overall, the geographic distribution of encounter potential was very dependent on both the behavioural ecology of the relevant wild mammal and the specific livelihood-related activities undertaken by people across the study area, with both location and landscape structure shaping where encounter potential was concentrated.

Elephant accounted for by far the greatest apparent potential for dangerous encounters with humans, consistent with existing evidence that they dominate HWC across many parts of Africa because of their size, intelligence and propensity for crop-raiding (20,53). Regarding activity overlap with all humans, the fishing camps along the Kilombero river in the northern of the WMA exhibited the greatest encounter potential, reflecting the role of these locations in supporting livelihoods for both resident custodian communities (19,34) and the traders who connect them to markets and supplies outside the WMA. While each of these regulated settlements may be home to a hundred or more people who store and sometimes even grow food crops at garden scale, it is also noteworthy that dozens of traders regularly visit them by canoe, bicycle or motorbike to buy their fish and/or sell them supplies. These camps are all situated amidst evergreen groundwater forest that provides suitable cover, food and water for elephant while also limiting visibility for people travelling along winding paths through the same dense vegetation at considerable speed (54). This interpretation is consistent with previous studies showing that elephant, bulls in particular, often remain close to habitat edges to access anthropogenic food sources (46), and that habitat use and human-elephant interactions are often concentrated in areas close to natural habitat, forest fringes, reserve boundaries and water sources (55,56).

However, regarding encounter potential for the subset of people practicing unauthorized livelihoods within the WMA and NNP, proximity to the settled-natural ecotone was the strongest predictor of risk. This reflects the fact that such covert activities not only make them especially vulnerable (Fig 2) they also often place them right within or beside well-used elephant habitat and transit pathways. As a result, this ecotone functions as a key zone of high potential risk, where elephant and particularly vulnerable people are most likely to come into contact. Furthermore, the inherent dangers of this apparent conflict zone may be exacerbated by attempts to ward off crop-raiding elephants that can increase aggressive behaviours, often culminating in injury or death (57–59).

Humans activity overlap with buffalo was more evenly distributed across natural areas and the peripheral settlements closest to them. These natural areas in ILUMA WMA are largely dominated by miombo woodland providing suitable habitat for buffalo (60). As for elephant, much of the human potential to encounter buffalo in these natural areas was associated with conducting unauthorized natural resource extraction activities that render them particularly vulnerable (Fig 2), such as charcoal burning, timber harvesting or meat poaching, suggesting that limited capacity of the WMA to enforce conservation regulations results in substantial amounts of otherwise preventable risk (19,34). De facto risk at unauthorized peripheral settlements within the WMA may also be exacerbated by their frequent establishment close to water sources: during the dry season, buffalo can exhibit reduced avoidance of cattle because water availability becomes a limiting resource, increasing overlap with human activities (61,62). However, regarding encounters with humans in general, including authorized WMA residents and visitors, fishing camps exhibited the highest encounter risk.

Consistent with the ecology of hippopotamus (63,64), potential for humans to encounter them was strongly tied to the availability of water of sufficient depth within ILUMA, at either the fishing camps along the Kilombero river or a small number of locations close to the Namkena river in the southeast. During the day, hippopotamus typically remain submerged or partially submerged in water to avoid heat stress (63,64). They emerge from water primarily at night to graze on terrestrial vegetation, often travelling several kilometres from their daytime resting sites (63,64). These substantial water sources are also centrally important to the livelihoods of people living nearby, be they fishers or farmers, respectively (65,66), so it is unsurprising that potential to encounter hippopotamus is highly focused on these areas.

Interestingly, potential for human encounter with hyena was greatest along the western boundary of ILUMA, where the WMA gives way to settlements and villages. Hyenas were also the only species for which appreciable encounter potential occurs in villages. This pattern is consistent with the known adaptability of hyena and their ability to exploit human-modified environments (67) by opportunistically scavenging anthropogenic food sources, such as waste dumps near settlements (68,69). Livestock, particularly goats and sheep, may provide an additional attractant in these areas, as their presence can bring hyena into close proximity to human settlements and increase encounter risk where livestock are kept (70). Hyenas exhibit primarily nocturnal (71) long distance foraging behaviours, increasing the likelihood of encounters with people at night. This is especially relevant for the most vulnerable people working in the WMA who sleep in poorly constructed makeshift shelters (Fig 2), a known risk factor for attack by another nocturnal large carnivore in Tanzania, specifically lion (72).

### 4.2 Study methodology limitations and advantages

Although this study provides useful insights into spatial distribution of where dangerous encounters with wildlife are most likely to occur, one particularly important limitation of the methodology applied should be given careful consideration when interpreting these results. Specifically, the overlap indices calculated herein are the products of largely indirect observations of the activities of human and wildlife, providing only relative indicators of potential for encounters rather than directly mapping actual conflict events. This is particularly important because human and wildlife activity may not coincide in exactly the same time and place within a given survey unit, especially where diurnal human activity overlaps with nocturnal wildlife activity, and vice versa. Therefore, these indices do not directly demonstrate where conflict actually occurred but instead identify areas where they are more likely, based on approximate co-occurrence of humans and dangerous large mammals, even though their periods of activity may only partially overlap (36).

Also, as explained in section 3.5, some underestimation of human activities that render them vulnerable to any dangerous wild mammals they encounter (Fig 2) may have occurred in the legally settled lands in the west of the study area because many of these were misclassified on the basis that these livelihood-related activities were usually perfectly legal outside of the WMA and NNP. Future studies designed with HWC risk mapping in mind should therefore use classification keys that record distinct activities and their legal status in that location as separate variables.

Another study limitation is that these surveys were not designed with this particular application in mind, and necessarily centres around perennial surface water bodies, so the purposive distribution of survey locations is neither random nor consistently systematic (37). While centring these activity surveys around year-round bodies of surface water inevitably introduces some biases, it also enhances sensitivity for the vast majority of wild mammals (37) and probably any humans who share those resources by choice or necessity (65,66).

Furthermore, this approach also has other important advantages. Since the overlap indices are based on large numbers of detectable activities that need to only approximately coincide with each other, rather than much rarer actual conflict events, they are likely much more statistically powerful for evaluating underlying determinants of hazard and risk. Also, unlike approaches that rely on injury, death or serious damage having already occurred, this approach allows risk to be assessed proactively without waiting for conflict events to take place. This also avoids the inherent limitations of “post mortem” conflict analyses, potentially enabling prospective assessment of preventable conflicts before sufficiently large numbers of such tragic events have been allowed to occur.

### 4.3 Potential for evidence-informed mitigation

With that in mind, it is important that this evidence is used to prospectively inform potential mitigation strategies despite obvious limitations of this indirect approach and the design of the studies that generated the data but was formulated for other purposes (33–38). In this study context, ecotonal boundaries between natural habitats and human settled areas within the ILUMA WMA represent priority locations targeted Village Game Scout (VGS) patrols, to prevent conflict with the two most important dangerous animals, namely elephant and buffalo, where it is currently most likely to occur. Similar approaches have demonstrated that the presence of rangers can reduce illegal activities like poaching in the surrounding locale, highlighting the potential value of strategically allocating patrol effort based on spatial patterns of risk (73). Of course, as this ecotone boundary between badly encroached and relatively well conserved natural habitat becomes better protected, it is likely to shift to the west and south, allowing elephant, buffalo and other wild animals to expand accordingly. Over the longer term, it is therefore important that such patrols are sustained across the entire WMA, and the greatest barrier to achieving that goal is simply resourcing those conservation enforcement activities.

More broadly, however, the bigger underlying issues for this WMA (18,19) and many others across Tanzania (74,75) are inadequate income as things stand and also a limited repertoire of financially viable business models with which to address that deficit. Having said that, recent developments within ILUMA demonstrate important progress towards strengthening the operational and financial sustainability of the WMA. In recent years, research activities undertaken across ILUMA (18,19,33–38) have provided a platform for sustained WMA-wide monitoring and patrol coverage, with Village Game Scouts (VGS) accompanying survey teams throughout extensive areas of the landscape. In addition, the establishment of a hunting company contract has created a new source of revenue for the WMA, while a recent $1,000,000 investment commitment from Reforest Africa provides further opportunities to support conservation and sustainable development initiatives. Furthermore, it may be possible to further develop the custodian community model exemplified by the authorized fishing camps in ILUMA (19,34) to other sustainable natural resource harvesting practices, such as seasonal grazing or silage harvesting, so that the resident monitoring functions these regulated settlements perform can be extended southwards all across the WMA (19).

Taking the existing fishing camps already established within ILUMA as an excellent example of resident custodian communities living within conservation areas, protective fencing may represent a useful intervention at such focal but clearly hazardous locations identified by this study, where appreciable numbers of elephant, buffalo, hippopotamus and hyena all overlap heavily with indispensable livelihood activities for local stakeholder communities of the WMA. Note, however, that mitigation tactics that combine physical barriers with community involvement may provide more sustainable alternatives to conventional fencing as a stand-alone intervention (76). More broadly, such mitigation efforts should combine targeted monitoring, community engagement and improved management of high-risk interface areas (77,78).

Beyond the more abundant elephant and buffalo, mitigation of human conflict with hippopotamus remains especially challenging, with relatively few studies experimentally evaluating the effectiveness of different approaches (79). Nevertheless, physical barriers like fences, cables and trenches have shown potential for preventing crop raiding, although their effectiveness is often constrained by high costs, maintenance requirements and the potential for habituation (79). Similarly, deterrent approaches such as lighting, acoustic disturbance and guarding have been used to discourage hippopotamus movements into agricultural areas, although many require continued human presence and may lose effectiveness over time (56,79). Given that most human–hippopotamus conflict occurs in or near water sources, habitat-based approaches like riparian buffer zones (strips of natural vegetation maintained along rivers to separate aquatic habitat from human land use) may provide a more sustainable solution by maintaining grazing areas for hippopotamus while reducing overlap between wildlife and human activities (79).

For hyena, mitigation presents different challenges due to their adaptability and association with human-modified environments. Since encounter risk was concentrated near villages and peripheral settlements, interventions should address the issues that attract them into human settlements by, for example, improving disposal of anthropogenic food waste and livestock carcasses (80). Regarding the hazards of living conditions that leave people highly vulnerable to hyena attack while asleep at night (Fig 2), the best way to mitigate those risks is simply to rigorously discourage encroachment upon the WMA to practice the unauthorized natural resource harvesting activities they are so closely associated with. That in turn means more patrols, improved spatial coverage with focal, well-regulated custodian communities (19,34), and ultimately better resourcing of the WMA through new business models that address current concerns of local stakeholder communities (18).

## Conclusion

Despite its limitations, this study nevertheless demonstrates that mapping potential encounter risk based on the spatial and temporal overlap of largely indirect activity indicators like tracks, spoor and signs appears to provide a proactive, pre-emptive and statistically powerful way to identify where hazardous encounters are most likely to occur, potentially offering considerable advantages over retrospective assessments of much sparser observations of actual realised conflict event after they have already occurred. While potential for encountering the two most apparently important dangerous animals, namely elephant and buffalo was closely associated with authorized fishing camps within the ILUMA WMA and the ecotone between substantially disturbed and largely intact natural land cover, the greatest potential for encountering hippopotamus occurred largely around two river beds, while hyena encounter potential was more closely associated with established villages and converted lands close to the western boundary of the ILUMA WMA. These new maps of hazard and risk suggest that effective HWC mitigation in this particular WMA is unlikely to arise from any single stand-alone intervention, but from targeted actions that reduce spatiotemporal overlap between people and wildlife by keeping them apart and/or protecting them against each other. By identifying where and when these overlaps occur, this new spatial risk mapping approach provides a practical framework for shifting management from reactive responses after conflict occurs towards proactive prevention.

## Author contributions

This study was planned by Gerry F. Killeen, Lucia J. Tarimo, Deogratius R. Kavishe, Felister Mombo and Ramiro D Crego and then implemented by Jude O’Regan with assistance from Lucia J. Tarimo, Gerry F. Killeen, Felister Mombo, Fidelma Butler, Ramiro Crego, Aristides Litaka, Elisa Manase and Mohamedi Kamuna. Jude O’Regan wrote the initial draft of the manuscript, which was then edited by Gerry F. Killeen in consultation with Lucia Tarimo, Ramiro Crego, Deogratius Kavishe and Felister Mombo. All authors read and commented on the manuscript and approved the final version for submission.

## Supporting information

O'Regan et al-Data on human-wildlife conflict in southern Tanzania

## Acknowledgements

The authors wish to thank the Village Game Scouts of ILUMA WMA, for their hard work and participation in field activities. We also thank all the governance, management and stakeholder communities of the ILUMA WMA for all their collaboration and kind assistance over the course of the study. Furthermore, we thank Dr. Emmanuel Kaindoa, Mr. Frederic Masanja and Mr Fadhili Sango for all the essential institutional support provided by the Ifakara Health Institute over the course of the study.

## Data availability statement

All the detailed quantitative human activity, animal activity and land cover-data data used for the analyses presented herein are freely available in the following supplementary file: https://doi.org/10.5281/zenodo.23195146

